# A novel biosynthetic gene cluster across the *Pantoea* species complex is important for pathogenicity in onion

**DOI:** 10.1101/2022.03.03.482764

**Authors:** Mei Zhao, Shaun Stice, Gi Yoon Shin, Teresa Coutinho, Ron Gitaitis, Brian Kvitko, Bhabesh Dutta

## Abstract

Onion center rot is caused by at least four species of *Pantoea* (*P. ananatis, P. agglomerans, P. allii, and P. stewartii* subsp. *indologenes*). Critical onion pathogenicity determinants for *P. ananatis* were recently described but whether those determinants are common among other onion-pathogenic *Pantoea* species remains unknown. In this work, we report onion pathogenicity determinants in *P. stewartii* subsp. *indologenes* and *P. allii*. We identified two distinct secondary metabolite biosynthetic gene clusters in different strains of onion pathogenic *P. stewartii* subsp. *indologenes*. One cluster is similar to the previously described HiVir phosphonate biosynthetic cluster of *P. ananatis* and another is a novel putative phosphonate biosynthetic gene cluster, which we name “Halophos”. The Halophos gene cluster was also identified in *P. allii* strains. Both clusters are predicted to be phosphonate biosynthetic clusters based on the presence of a characteristic phosphenolpyruvate phosphomutase (*pepM*) gene. The deletion of *pepM* gene from either the *P. stewartii* subsp. *indologenes* HiVir or Halophos clusters caused loss of necrosis on onion leaves and red onion scales, and resulted in significantly lower bacterial populations compared to the corresponding wildtype and complemented strains. Seven (*halB*-*halH*) out of eleven genes (*halA*-*halK*) in the Halophos gene cluster are required for onion necrosis phenotypes. The onion non-pathogenic strain PNA15-2 gained the capacity to cause foliar necrosis on onion via exogenous expression of a minimal seven gene Halophos cluster (*halB -halH)*. Furthermore, cell-free culture filtrates of PNA14-12 expressing the intact Halophos-gene cluster caused necrosis on onion leaves consistent with the presence of a secreted toxin. Together, these observations indicated that *pepM* genes in both phosphonate biosynthetic gene clusters (HiVir and Halophos) are important for *Pantoea* spp. onion pathogenicity and the biosynthetic product of the Halophos cluster causes necrosis on onion leaf tissue. Overall, this is the first report of onion pathogenicity determinants in *P. stewartii* subsp. *indologenes* and *P. allii*.

**Author summary:** Onion center rot is caused by multiple *Pantoea* species including *P. stewartii* subsp. *indologenes* and *P. allii*. We identified two distinct secondary metabolite biosynthetic clusters associated with onion pathogenic strains, the validated HiVir phosphonate cluster and a putative phosphonate biosynthetic cluster that we named as Halophos based on the associated “halo” phenotype on the red onion scales. We found that *pepM* genes from each cluster (HiVir and Halophos) are required for onion infection by *P. stewartii* subsp. *indologenes* and *P. allii* but not for millet infection by *P. stewartii* subsp. *indologenes*. Conversely, the T3SS was important for millet infection by *P. stewartii* subsp. *indologenes* but not onion infection. Induction of the intact Halophos cluster was associated with the accumulation of a necrosis-inducing factor in culture, which suggests it might be a secreted phytotoxin. Seven of the eleven Halophos cluster genes are required for onion necrosis phenotypes and expression of this minimal cluster conferred a limited onion necrosis phenotype to an onion-non-pathogenic *Pantoea* strain. We provide evidence of a Halophos biosynthetic gene cluster to be associated with onion pathogenicity in strains of *P. stewartii* subsp. *indologenes* and *P. allii*.

## Introduction

Bacteria in the genus *Pantoea* are ubiquitous and form a wide range of interactions with eukaryotic hosts including plants, fungi, insects, and humans [1]. Strains of at least four *Pantoea* species, *P. ananatis* [2], *P. agglomerans* [3, 4], *P. allii* [4], and *P. stewartii* subsp. *indologenes* [5], have been associated with onion center rot. Onion center rot can cause severe yield losses both in the field and in storage, and in some cases, economic losses up to 90% have been experienced [2]. In addition, recently, *P. dispersa* was reported to cause bulb decay of onion [6]. Commercial onion cultivars with resistance to *Pantoea* spp. have not been identified [7]. Thus, there is a need to understand pathogenesis mechanisms in *Pantoea* spp. that may potentially provide necessary information for future breeding efforts.

Two virulence factors that distinguish between onion-pathogenic and non-pathogenic strains of *P. ananatis* have recently been described [8]. Most bacterial pathogens depend on specialized virulence protein based secretion systems for pathogenicity [9]. Specifically, the majority of gram-negative bacterial plant pathogens are dependent on either a virulence-associated Hrp type III secretion system (T3SS) to deliver immune-dampening effector proteins or the type II secretion system (T2SS) to deliver the plant cell wall degrading enzymes associated with soft rot diseases [9]. However, *P. ananatis* lacks type II and type III secretion systems [10]. Instead, *P. ananatis* requires a HiVir (<u>Hi</u>gh <u>Vir</u>ulence) gene cluster for pathogenicity on onion [8]. Disruption of the HiVir cluster in *P. ananatis* resulted in the loss of pathogenicity on onion foliage, on scales of red onion bulbs, and on whole onion bulbs [7, 8, 11]. The HiVir gene cluster was predicted to encode synthesis of a phosphonate compound based on the presence of a characteristic *pepM*, phosphoenolpyruvate (PEP) phosphonomutase gene within the cluster [8] and the novel phosphonate compound designated pantaphos (2-(hydroxy[phosphono]methyl)maleate) was shown to be synthesized by the HiVir cluster and cause extreme tissue damage in onion bulbs [11].

*Pantoea stewartii* subsp. *indologenes* causes leaf spots on foxtail millet and pearl millet [12]. It was recently reported that some strains were also pathogenic on onion in Georgia, USA [5]. The epithet *P. stewartii* subsp. *indologenes* pv. *cepacicola* was proposed for strains (PNA03-3, PNA14-9, PNA14-12 and PNA14-11) based on the host range tests on *Allium* spp., including onion and also on foxtail millet, pearl millet, and oat [13]. Interestingly, the HiVir gene cluster was identified in *P. stewartii* subsp. *indologenes* strain PNA03-3 but absent in PNA14-9, PNA14-11, and PNA14-12 based on a PCR assay with HiVir-specific primers [13]. These observations were intriguing as despite the absence of the HiVir cluster these strains were pathogenic on onion. We hypothesized that the HiVir gene cluster was important for PNA03-3 onion pathogenicity and that other pathogenicity factors could potentially be involved in onion pathogenicity in strains, PNA14-9, PNA14-11, and PNA14-12. Based on the annotation of secondary metabolite biosynthetic gene clusters using antiSMASH 6.0 [14], we found that *P. stewartii* subsp. *indologenes* strain PNA14-12 as well as *P. allii* type strain LMG24248^T^ were predicted to possess a completely distinct putative phosphonate biosynthetic cluster. The strains that possess this putative gene cluster produced pink halo around the necrotic tissue on red-onion scales and hence a name “Halophos” is being proposed here. We hypothesized that the Halophos gene cluster would be important in *P. stewartii* subsp. *indologenes* and *P. allii* for pathogenicity in onion. Hence, here we provided evidence of the role of *pepM* from Halophos in onion pathogenicity in *P*. *stewartii* subsp. *indologenes* and *P. allii*. In addition, we also characterized the role of *pepM* from the HiVir gene cluster in *P. stewartii* subsp. *indologenes* strain PNA03-3. Since *P. stewartii* subsp. *indologenes* also encodes T3SS, which displayed the closest similarity to *P. stewartii* subsp. *stewartii*, we also determined the role of the T3SS for *P. stewartii* subsp. *indologenes* pathogenicity on onion and pearl millet.

## Results

### The presence of phosphonate biosynthetic gene clusters correlated with *Pantoea stewartii* subsp. *indologenes* pathogenicity in onion

In order to understand the onion pathogenicity mechanisms of *P. stewartii subsp. indologenes*, the genomes (under the bio-project PRJNA676043) of onion-pathogenic strains (*n*=4, PNA03-3, PNA14-9, PNA14-11, and PNA14-12) and onion-non-pathogenic strains (*n*=13, PNA15-2, PANS07-4, PANS07-6, PANS07-10, PANS07-12, PANS07-14, PANS99-15, NCPPB1562, NCPPB1877, NCPPB2275, NCPPB2281, NCPPB2282, and LMG2632) reported in Koirala *et al*. (2021) [13] were analyzed for the presence of the HiVir and Halophos gene cluster sequences. Surprisingly, the phosphonate biosynthetic gene cluster HiVir was found only in PNA03-3. The Halophos gene cluster including a PepM homolog was identified in onion pathogenic strains PNA14-9, PNA14-11, and PNA14-12. The phosphonate biosynthetic gene clusters (HiVir or Halophos) were present only in onion pathogenic strains (PNA03-3, PNA14-9, PNA14-11, and PNA14-12), suggesting that the putative phosphonate compounds may be essential for pathogenicity in onion. The Halophos cluster is adjacent to a phage/plasmid primase gene in *P*. *stewartii* subsp. *indologenes* PNA14-9, PNA14-11, and PNA14-12. The Halophos cluster from PNA14-12 is located within a genomic island predicted by IslandViewer 4. In addition, the Halophos cluster has a lower GC%, higher average effective number of codons (Nc), and lower average codon adaptation index (CAI), compared to their corresponding whole genomes (Table 1), altogether suggesting acquisition through horizontal gene transfer (HGT).

**Table 1.** Sequence signatures of Halophos gene cluster and PNA14-12 genome.

| Feature | Halophos gene cluster | PNA14-12 genome |
| --- | --- | --- |
| Average ORF GC content (%) | 46.5 ± 2.9 | 53.5 ± 5.6 |
| Average effective number of codons (Nc) | 56.09 ± 2.39 | 46.36 ± 5.88 |
| Average codon adaptation index (CAI) | 0.25 ± 0.03 | 0.31 ± 0.08 |

± standard deviation.

The Halophos gene cluster is predicted to contain eleven co-transcribed genes (Table 2), which we have named *halA* through *halK*, encoding one iron-containing alcohol dehydrogenase (*halA*), alcohol dehydrogenase (*halB*), N-acetyl-gamma-glutamyl-phosphate reductase (*halC*), phosphoglycerate dehydrogenase (*halD*), phosphoenolpyruvate mutase (*halE*), FAD-NAD(P)-binding protein (*halF*), aminotransferase (*halG*), phosphoglycerate kinase (*halH*), MFS transporter (*halI*), AMP-binding enzyme (*halJ*), and pyridoxamine 5’-phosphate oxidase (*halK*) (Fig 1 and Table 2). Notably, the *halJ* sequence is predicted to possess an adenylation domain and a condensation domain, suggesting that it might be involved in non-ribosomal peptide synthetase activity.

**Table 2.**
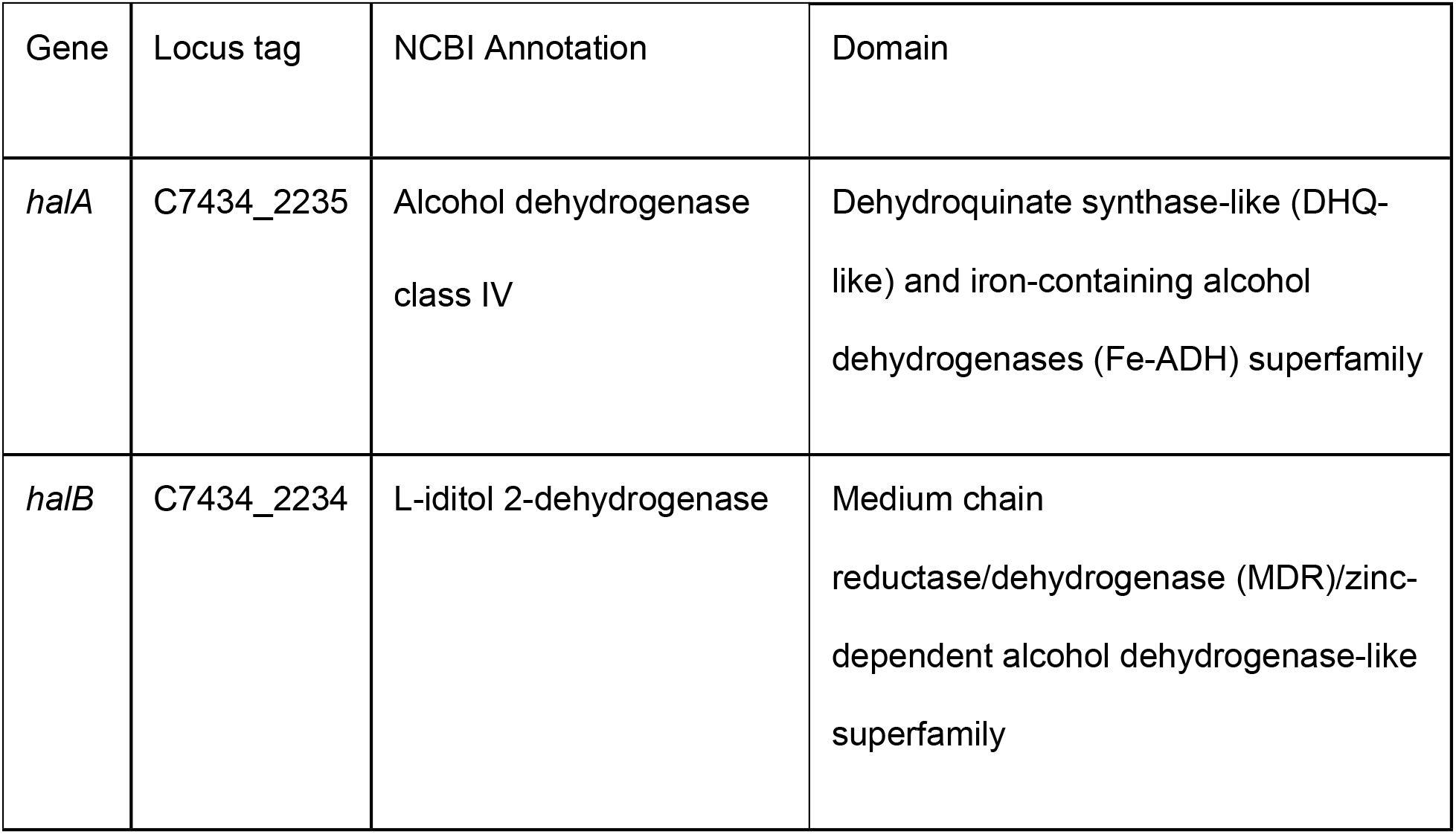
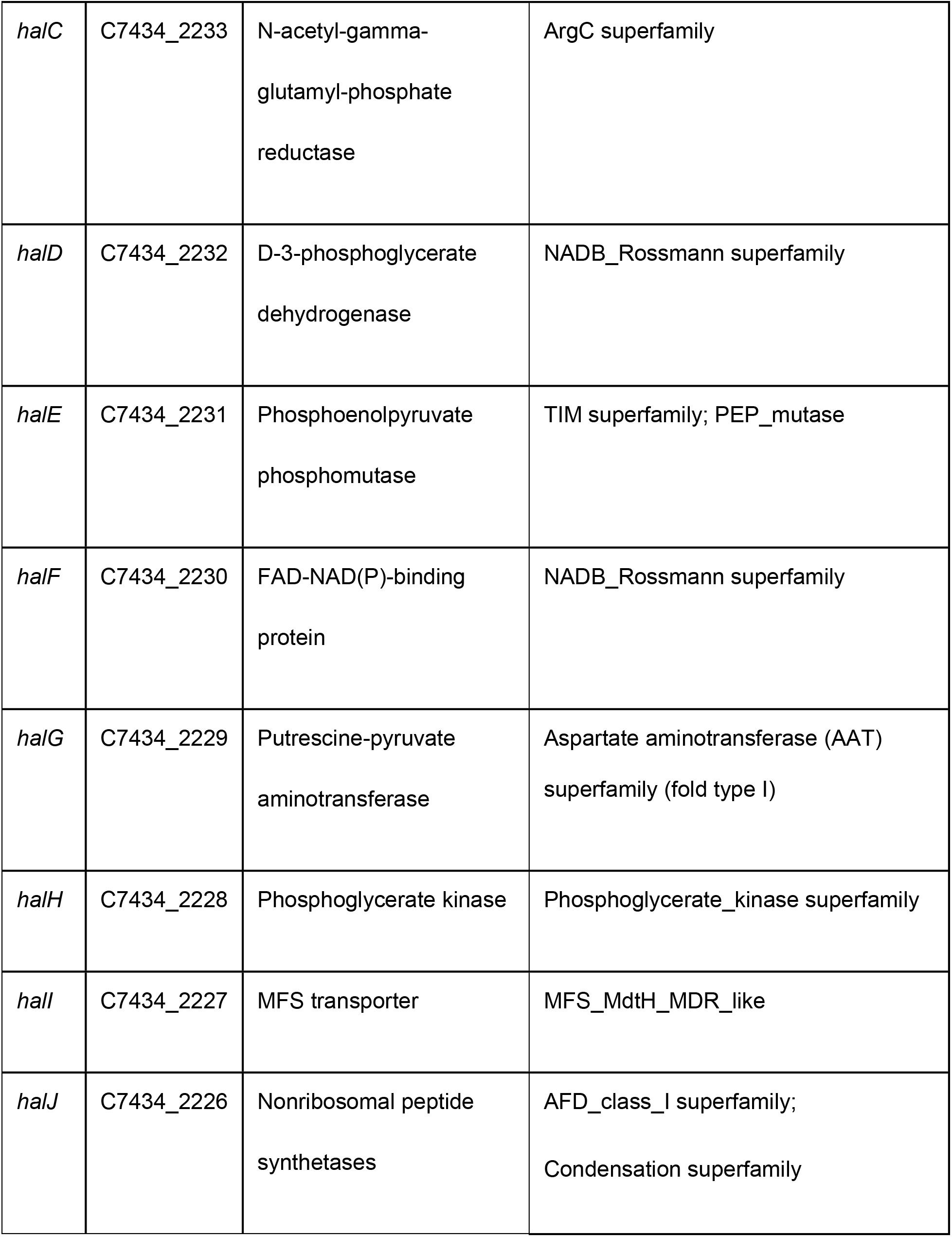
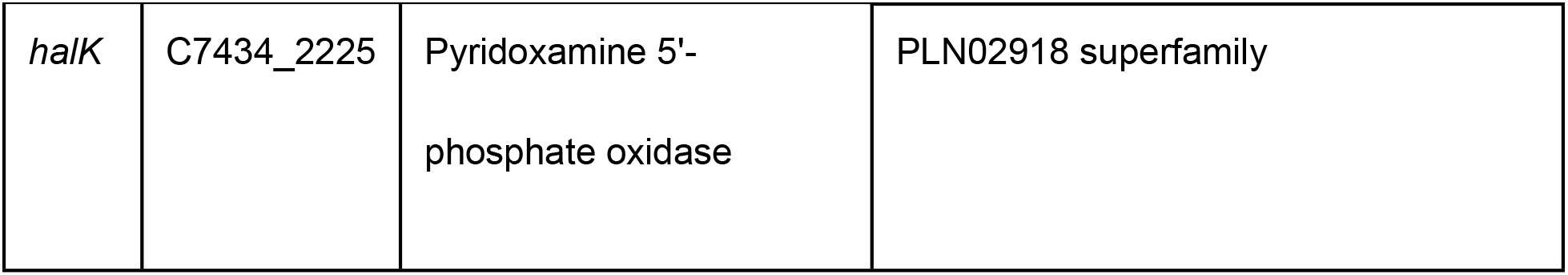
Genes in the Halophos cluster of PNA14-12. Gene, proposed gene name; Locus tag, NCBI locus tag from accession NZ_SOAJ01000001 for PNA14-12; Domain, conserved domains within the protein sequences obtained from https://www.ncbi.nlm.nih.gov/Structure/cdd/wrpsb.cgi.

**Fig 1.**
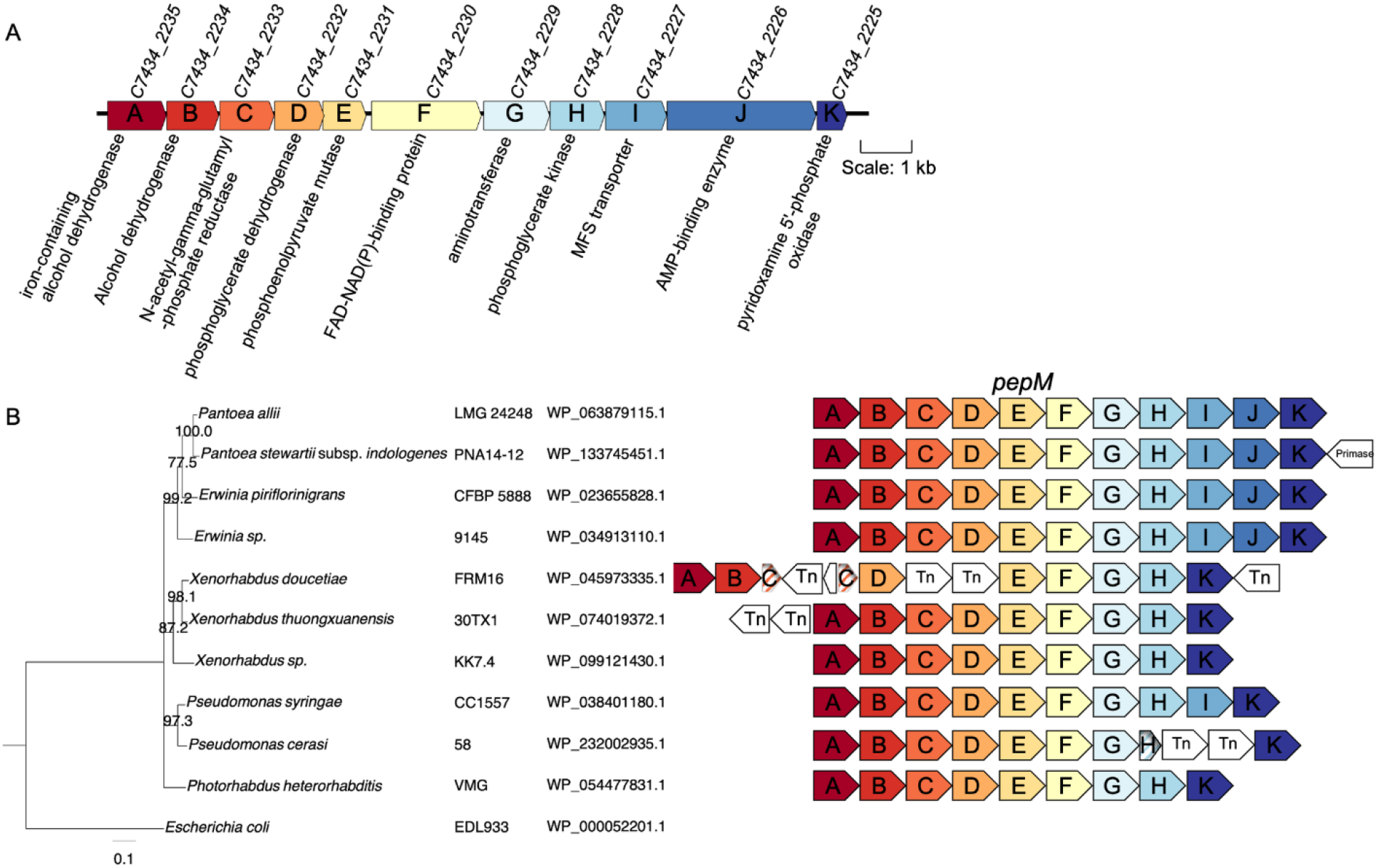
Halophos gene cluster structure and synteny. (A) Diagram of the Halophos gene cluster of *Pantoea stewartii* subsp. *indologenes* PNA14-12. The gene locus accessions are according to PNA14-12 annotation (accession NZ_SOAJ01000001). The scale represents 1 kb. (B) A phylogenetic tree based on selected PepM protein sequences of organisms that have Halophos-like gene clusters. Ten PepM protein sequences and *Escherichia coli* 2-methylisocitrate lyase sequence (as an outgroup) were used for multiple sequence alignment using MAFFT. The alignment was trimmed to the same length and used for constructing a neighbor-joining tree. The bootstrap support values > 70% were shown at the node. The conserved clusters (not to scale) were colored with the same colors representing homologous proteins. Non-conserved proteins were in white, and fragmented pseudogenes were in stripes. Transposase genes were labeled as Tn.

### Bioinformatics analysis of the Halophos-like gene clusters

Analysis with antiSMASH predicted that the Halophos gene clusters of PNA14-12 and *P. allii* LMG24248 encode phosphonate secondary metabolite gene clusters. Using PNA14-12 Halophos nucleotide and protein sequences for blastn and tblastn searches, we identified 34 strains that encoded Halophos-like gene clusters (S1 Table). The strains belonged to the genera *Pantoea, Erwinia, Pseudomonas, Xenorhabdus*, and *Photorhabdus*. *Pantoea allii* strains shared 85% sequence identity with the PNA14-12 Halophos nucleotide sequence. *Erwinia* strains shared 68% to 71% sequence identity with PNA14-12 Halophos nucleotide sequence. *Xenorhabdus, Photorhabdus* and *Pseudomonas spp*. had 65%-72% nucleotide sequence identity with the PNA14-12 Halophos sequence (at a sequence coverage of 30%-46%) (S1 Table). Except for *Pseudomonas syringae* CC1557 (isolated from snow), all other strains that encoded Halophos-like sequences were isolated from plants or nematodes. Among them, *Pantoea* and *Erwinia* strains showed the same gene synteny. *Xenorhabdus* and *Photorhabdus* strains do not have *halI* (MFS transporter) and *halJ* (AMP-binding enzyme) in their Halophos-like gene clusters. The *halC* gene in *X. doucetiae* strain FRM16 was disrupted by a transposase gene. Also, *Pseudomonas* strains do not have *halJ* in their gene clusters and *P. cerasi* strains also have fragmented *halH* (Fig 1B).

Comparison of the two phosphonate biosynthetic gene clusters, HiVir and Halophos revealed that only two genes were similar based on predicted annotations: *pepM* and the major facilitator superfamily (MFS) transporter gene (Tables 2 and 3). These *pepM* genes shared 35.1% amino acid identity. Both PepM proteins (HalE from Halophos and HvrA from HiVir) possessed the PEP mutase domain (Tables 2 and 3) and the key PepM catalytic motif (EDKXXXXXNS). Similarly, the MFS transporter genes from the two phosphonate clusters shared 32.1% sequence identity. The MFS transporters from the HiVir and the Halophos gene clusters belonged to the macrolide efflux protein A (MefA) family and the multidrug resistance protein MdtH family, respectively. These findings suggest their involvement in transporting different molecules.

**Table 3.** Genes in the HiVir cluster of PNA03-3. Gene, proposed gene name; Locus tag, NCBI locus tag from accession NZ_QICO01000001 for PNA03-3; Domain, conserved domains within the protein sequences obtained from https://www.ncbi.nlm.nih.gov/Structure/cdd/wrpsb.cgi.

| Gene | Locus tag | NCBI Annotation | Domain |
| --- | --- | --- | --- |
| *hvrA* | C7433_102431 | Phosphoenolpyruvate<br>phosphomutase | TIM superfamily; PEP_mutase |
| *hvrB* | C7433_102432 | LLM class flavin-dependent<br>oxidoreductase | Flavin-utilizing monooxygenases<br>superfamily |
| *hvrC* | C7433_102433 | Homocitrate synthase NifV | aksA superfamily |
| *hvrD* | C7433_102434 | 3-isopropylmalate<br>dehydratase large subunit | Aconitase superfamily |
| *hvrE* | C7433_102435 | 3-isopropylmalate<br>dehydratase small subunit | Aconitase_swivel superfamily |
| *hvrF* | C7433_102436 | Class I SAM-dependent<br>methyltransferase | AdoMet-MTases superfamily |
| *hvrG* | C7433_102437 | GNAT family N-acetyltransferase | NAT_SF superfamily |
| *hvrH* | C7433_102438 | Hypothetical protein | PRK12767 superfamily (carbamoyl<br>phosphate synthase-like protein) |
| *hvrI* | C7433_102439 | MFS transporter | MFS_MefA_like |
| *hvrJ* | C7433_102440 | Hypothetical protein | No putative conserved domains |
| *hvrK* | C7433_102441 | Flavin reductase | PNPOx/FlaRed_like superfamily |

### The HiVir or Halophos gene cluster is critical for onion pathogenicity in *P. stewartii* subsp. *indologenes* but the T3SS is not

The *pepM* gene is present in both phosphonate biosynthetic clusters (HiVir and Halophos) in *P. stewartii* subsp. *indologenes*. In order to decipher the role of *pepM* genes in HiVir and Halophos clusters, the *pepM* gene was deleted in these two distinct phosphonate clusters in *P. stewartii subsp. indologenes* strains (PNA03-3 and PNA14-12).

*Pantoea stewartii* subsp. *indologenes* PNA03-3Δ*pepM* reached significantly lower population levels than PNA03-3 wildtype (*P* < 0.0001) (Fig 2A) when inoculated into red onion scales in addition to losing the associated red scale necrosis phenotype. Both bacterial load and necrosis phenotypes were complemented by the expression of the Halophos *pepM* gene (*halE*) from PNA14-12. Complementation of the PNA03-3 HiVir cluster *pepM* deletion with the *pepM* gene from PNA14-12 provides genetic validation that HalE is a functional PepM enzyme.

**Fig 2.**
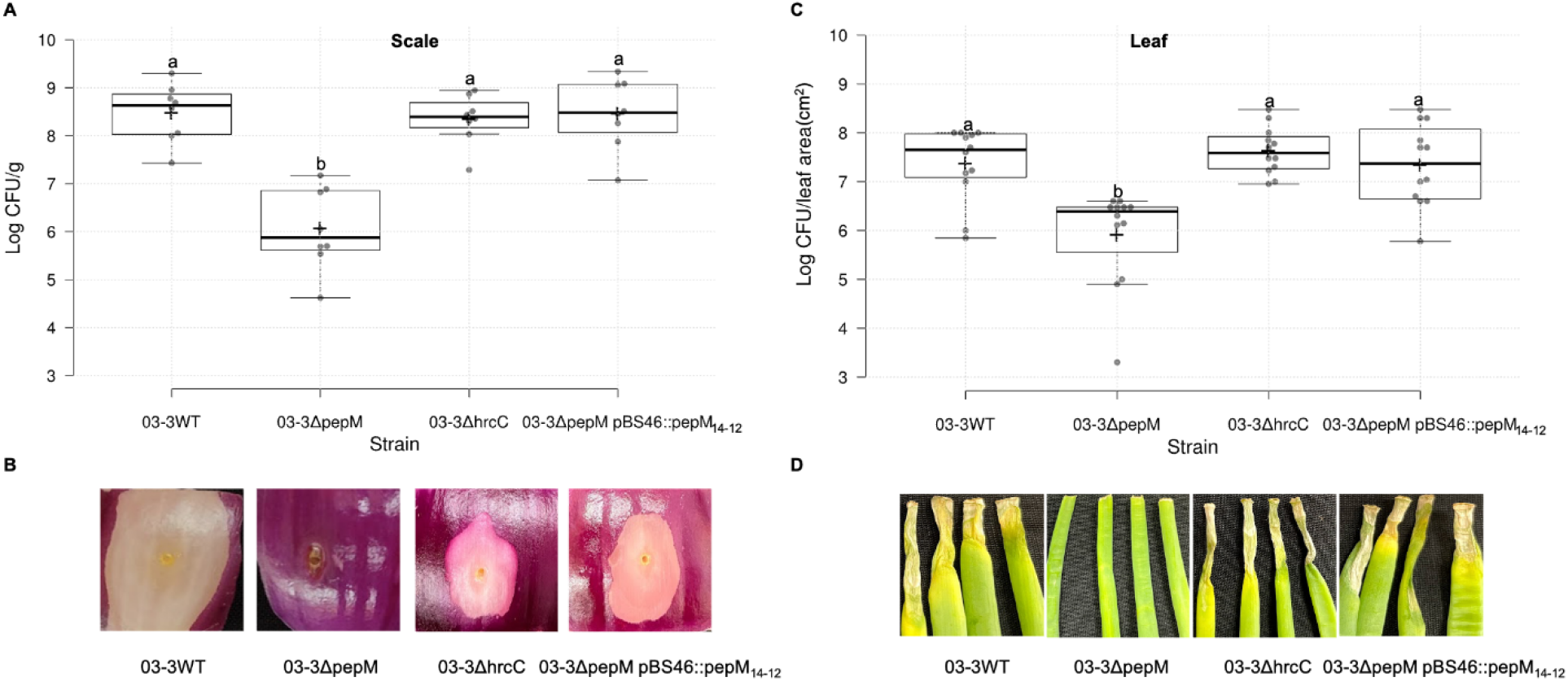
*pepM* was vital for *Pantoea stewartii* subsp. *indologenes* PNA03-3 pathogenicity on onion scale and leaf. Bacterial populations in onion scale (A) and leaf (C) tissues inoculated with *P. stewartii* subsp. *indologenes* PNA03-3 wildtype and mutants and representative symptoms produced on red onion scales (cv. Red Barret) (B) and 6-week-old onion leaves (cv. Century) (D). PNA03-3 wildtype (WT), 03-3*ΔpepM* mutant, 03-3*ΔhrcC* mutant, and 03-3Δ*pepM* pBS46∷*pepM_14-12_* complement strains were inoculated onto red onion scale and leaves at 10^4^ CFU (*n* = 4) for A, C, and D, and 10^6^ CFU for B. Samples and images were taken at 4 days post inoculation (dpi). For onion scale necrosis assay, tissue samples (0.2 cm x 0.2 cm) were taken 0.5 cm away from the inoculation point, weighed, and macerated in sterilized H_2_O, and plated on LB agar with rifampicin. Colonies were counted 24 h after incubation and converted to Log_10_ CFU/g. For onion leaf inoculation assays, samples (0.5 cm long) were taken 0.3 cm away from the inoculation point, processed similarly to the scale samples, and the colonies were enumerated as Log_10_ CFU/leaf area (cm^2^). Center lines show the medians; box limits indicate the 25^th^ and 75^th^ percentiles; whiskers extend 1.5 times the interquartile range from the 25^th^ and 75^th^ percentiles; crosses represent sample means; data points are plotted as grey circles as determined by R software. The experiment was conducted at least twice and sample points *n* = 8 and 12 were shown for graph A and C, respectively. Different letters indicate significant differences (*P* = 0.05) among treatments according to Tukey-Kramer’s honestly significant difference test.

On onion leaves, *P. stewartii* subsp. *indologenes* PNA03-3Δ*pepM* reached significantly lower population levels than the PNA03-3 wildtype at 4 dpi (*P* < 0.0001) (Fig 2C). In terms of symptom severity, at 4 dpi, 03-3Δ*pepM* did not show typical necrosis on onion leaves (lesion length = 0 cm, Figs 2D and S1) and was complemented by *pepM_14-12_*. Interestingly inactivation of the T3SS by deletion of the *hrcC* gene did not result in phenotypic changes to population levels or onion disease symptoms compared to the wildtype strain.

Similar to the above observations on the red onion scale, *P. stewartii* subsp. *indologenes* PNA14-12Δ*pepM* reached significantly lower population levels than PNA14-12 wildtype (*P* < 0.0001) (Fig 3A). In terms of symptom development, at 4 dpi, 14-12WT, 14-12Δ*hrcC*, and 14-12Δ*pepM* pBS46∷*pepM_14-12_* showed distinct pink halo phenotypes, while 14-12Δ*pepM* did not display any symptoms on the red onion scale (Fig 3B).

**Fig 3.**
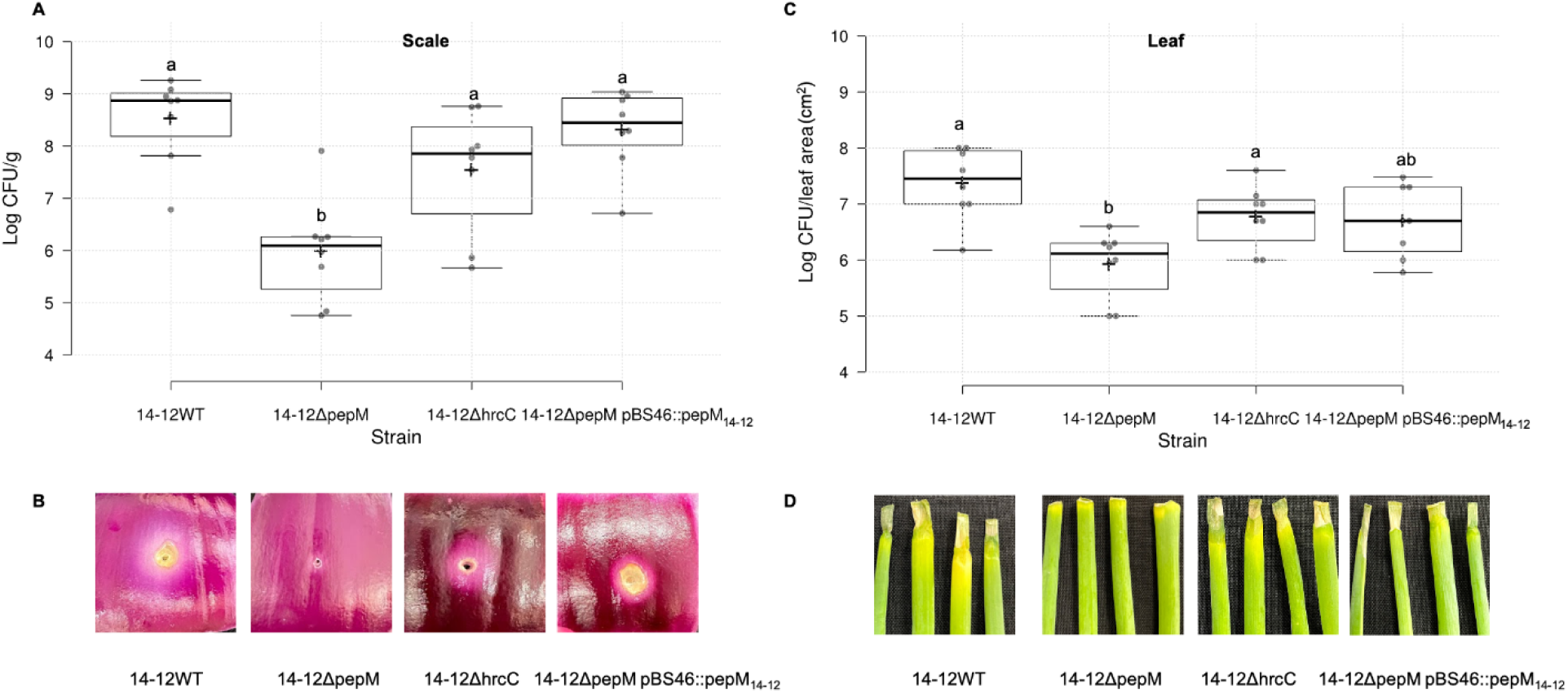
*pepM* was vital for *Pantoea stewartii* subsp. *indologenes* PNA14-12 pathogenicity on onion scale and leaf. Bacterial population levels in onion scale (A) and leaf (C) tissues inoculated with *P. stewartii* subsp. *indologenes* PNA14-12 wildtype and mutants, and representative symptoms produced on red onion scales (cv. Red Barret) (B) and 6-week-old onion leaves (cv. Century) (D). PNA14-12 wildtype (WT), 14-12*ΔpepM* mutant, 14-12*ΔhrcC* mutant, and 14-12Δ*pepM* pBS46∷*pepM_14-12_* complement strains were inoculated onto red onion scale and leaves at 10^4^ CFU (*n* = 4) for A, C, and D, and 10^6^ CFU for B. Samples and images were taken at 4 days post inoculation (dpi). For onion scale necrosis assay, tissue samples (0.2 cm x 0.2 cm) were taken 0.5 cm away from the inoculation point, weighed, and macerated in sterilized H_2_O and plated on LB agar with rifampicin. Colonies were counted 24 h after incubation and converted to Log_10_ CFU/g. For onion leaf inoculation assays, samples (0.5 cm long) were taken 0.3 cm away from the inoculation point, processed similarly to the scale samples, and the colonies were enumerated as Log_10_ CFU/leaf area (cm^2^). Center lines show the medians; box limits indicate the 25^th^ and 75^th^ percentiles; whiskers extend 1.5 times the interquartile range from the 25^th^ and 75^th^ percentiles; crosses represent sample means; data points are plotted as grey circles as determined by R software. The experiment was conducted twice and all sample points (*n* = 8) were shown. Different letters indicate significant differences (*P* = 0.05) among treatments according to Tukey-Kramer’s honestly significant difference test.

On onion leaves, *P. stewartii* subsp. *indologenes* PNA14-12Δ*pepM* reached significantly lower population levels than the PNA14-12 wildtype at 4 dpi (*P* = 0.0007) (Fig 3C). In terms of symptom severity, at 4 dpi, 14-12Δ*pepM* did not show typical necrosis on onion (lesion length = 0 cm, Figs 3D and S1), and was complemented by the *pepM_14-12_*. Similar to PNA03-3, inactivation of the T3SS by deletion of the *hrcC* gene in PNA14-12 did not result in phenotypic changes to population levels or onion disease symptoms compared to the wildtype strain.

Our initial nucleotide sequence similarity search of the Halophos gene cluster showed that another onion pathogen, *P. allii* LMG24248, also encodes the Halophos gene cluster (Fig 1 and S1 Table). In order to determine the role of *pepM* in *P. allii* LMG24248, the *pepM* gene of the Halophos was deleted. *Pantoea allii* LMG24248Δ*pepM* reached significantly lower population levels than LMG24248 wildtype (*P* < 0.0001) and did not display symptoms on onion scales and leaves (Fig 4). Both bacterial population levels and symptoms were complemented by the expression of the Halophos *pepM* gene (*halE*) from LMG24248.

**Fig 4.**
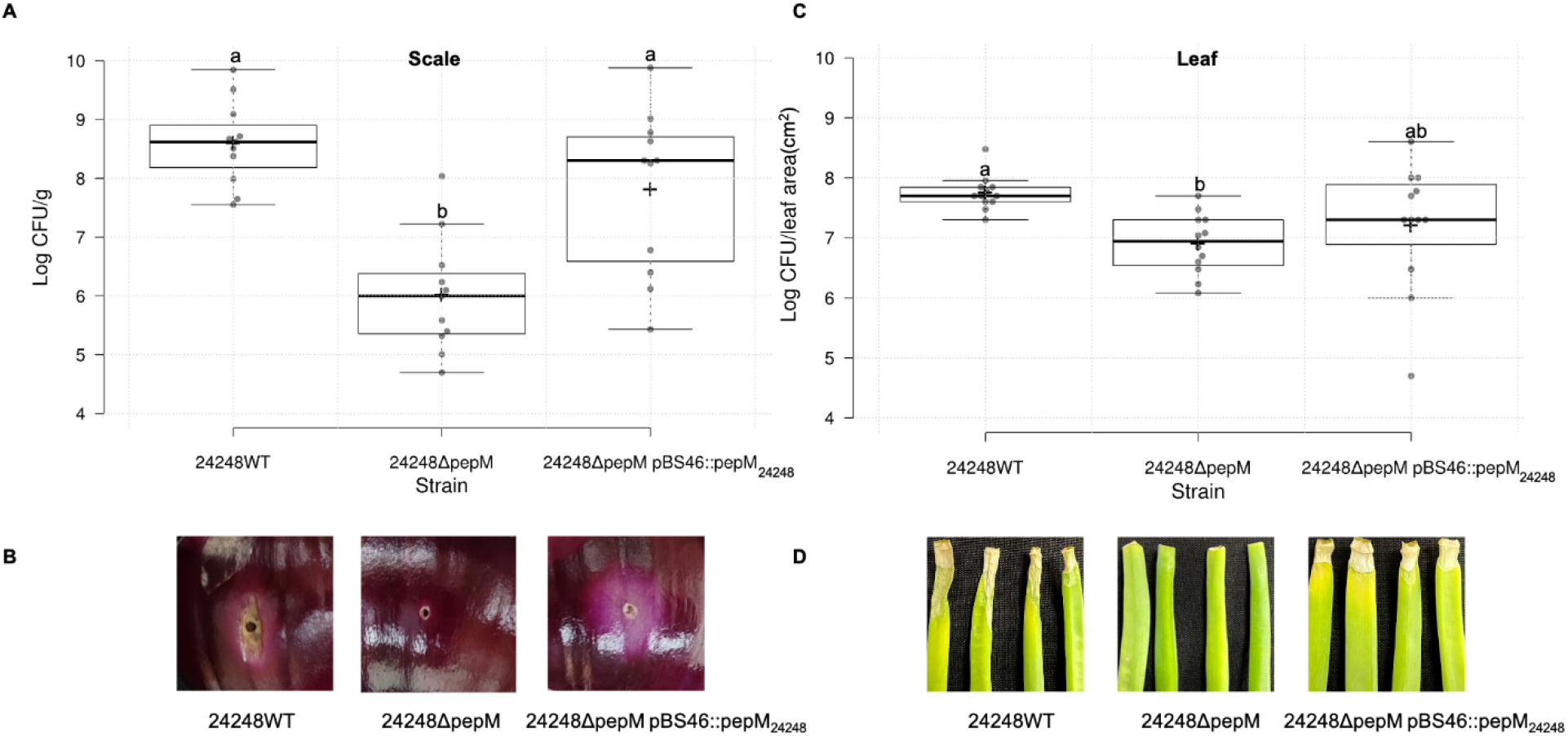
*pepM* was vital for *P. allii* LMG24248 pathogenicity on onion scale and leaf. Bacterial population levels in onion scale (A) and leaf (C) tissues inoculated with *P. allii* LMG24248 wildtype and mutants and representative symptoms produced on red onion scales (cv. Red Barret) (B) and 6-week-old onion leaves (cv. Century) (D). 24248 wildtype (WT), 24248*ΔpepM* mutant, and 24248Δ*pepM* pBS46∷*pepM_24248_* complement strains were inoculated onto red onion scale and leaves at 10^4^ CFU (*n* = 4) for A, C, and D, and 10^6^ CFU for B. Samples and images were taken at 4 days post inoculation (dpi). For onion scale necrosis assay, tissue samples (0.2 cm x 0.2 cm) were taken 0.5 cm away from the inoculation point, weighed, and macerated in sterilized H_2_O and plated on LB agar with rifampicin. Colonies were counted 24 h after incubation and converted to Log_10_ CFU/g. For onion leaf inoculation assays, samples (0.5 cm long) were taken 0.3 cm away from the inoculation point, processed similarly to the scale samples, and the colonies were enumerated as Log_10_ CFU/leaf area (cm^2^). Center lines show the medians; box limits indicate the 25^th^ and 75^th^ percentiles; whiskers extend 1.5 times the interquartile range from the 25^th^ and 75^th^ percentiles; crosses represent sample means; data points are plotted as grey circles as determined by R software. The experiment was conducted three times and all sample points (*n* = 12) were shown. Different letters indicate significant differences (*P* = 0.05) among treatments according to Tukey-Kramer’s honestly significant difference test.

We sequenced seven additional *P. allii* strains (BD381, BD382, BD383, BD386, BD387, BD388, and BD389) isolated from onion in South Africa. Based on whole genome sequencing, all seven *P. allii* strains encoded the Halophos-like gene cluster and their gene content and gene orientations within the cluster were identical. All seven strains caused foliar necrosis upon inoculation in onion leaf and displayed pink halo phenotypes in the red onion scale assays (S2 Fig).

### Contributions of the *hal* genes (Halophos) in *P. stewartii* subsp. *indologenes* PNA14-12 on onion pathogenicity

In order to determine the roles of other genes besides *pepM/halE* in the Halophos gene cluster, we made single mutants of each gene in PNA14-12, inoculated them on onion scale and leaves, and assessed the symptoms at 5 dpi. The strains PNA14-12WT, Δ*halA*, Δ*halI*, Δ*halJ*, and Δ*halK* showed distinct pink halo phenotypes on the red onion scales and caused necrosis on onion leaf, while Δ*halB*, Δ*halC*, Δ*halD*, Δ*halF*, and Δ*halG* did not display any symptoms on the red onion scale and on onion leaf (Figs 5A and 5B). The Δ*halH* deletion mutant showed no symptoms on the red onion scales but showed reduced foliar necrosis (Figs 5A and 5B). The complement strains of *halB, halC, halD, halF, halG*, and *halH* showed pink halo phenotypes and foliar necrosis (Figs 5A and 5B). In terms of disease severity on onion leaf, Δ*halB*, Δ*halC*, Δ*halD*, Δ*halF*, Δ*halG* displayed no lesion (lesion length = 0 cm, Fig 5C), while the mean lesion length for Δ*halH* was 1.0 cm, which was significantly lower than the 14-12WT (2.9 cm), Δ*halA* (2.1 cm), Δ*halI* (2.1 cm), Δ*halJ* (2.4 cm), and Δ*halK* (2.2 cm). The complement strains of *halBcomp* (1.0 cm), *halC*comp (1.1 cm), *halD*comp (2.5 cm), *halF*comp (1.0 cm), *halG*comp (2.3 cm), and *halH*comp (2.3 cm) showed significantly higher mean lesion length than that of their corresponding mutant strains (*P* < 0.0001) (Fig 5C). The segment of seven contiguous *hal* genes (*halB to halH*) was expressed in an onion-non-pathogenic *P. stewartii* subsp. *indologenes* strain PNA15-2. On onion leaf, the wildtype *P. stewartii* subsp. *indologenes* PNA15-2 (lack Halophos) did not cause foliar necrosis, while the transformant PNA15-2 pBS46∷*halB-H* induced limited necrotic lesions (mean length = 0.4 cm) (Figs 5D and 5E).

**Fig 5.**
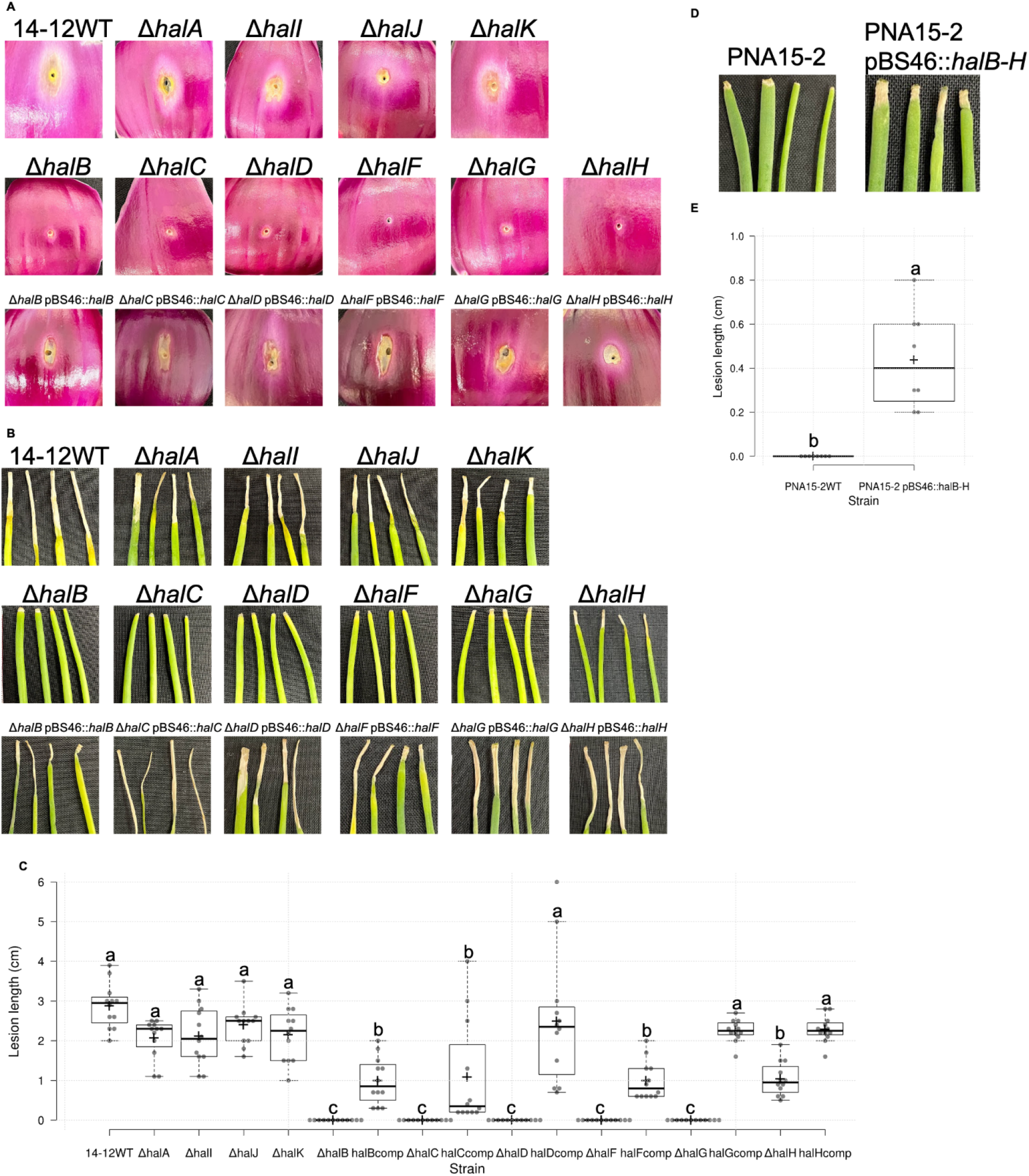
The *halB-H* genes are critical for *Pantoea stewartii* subsp. *indologenes* PNA14-12 pathogenicity on onion scale and leaf. Representative symptoms produced on red onion scales (cv. Red Barret) (A) and 6-week-old onion leaves (cv. Century) (B). PNA14-12 wildtype (WT), *hal* mutants, *hal* complement strains were inoculated onto red onion scale and leaves at 10^6^ CFU (*n* = 4). Samples and images were taken at 5 days post inoculation (dpi). (C) Lesion length on onion leaf. Complement strains are noted as halXcomp. Center lines show the medians; box limits indicate the 25^th^ and 75^th^ percentiles; whiskers extend 1.5 times the interquartile range from the 25^th^ and 75^th^ percentiles; crosses represent sample means; data points are plotted as grey circles as determined by R software. The experiment was conducted at least twice and all sample points were shown (*n* = 12 and 8 for graph C and E, respectively). Different letters indicate significant differences (*P* = 0.05) among treatments according to Tukey-Kramer’s honestly significant difference test.

### Bacterial culture filtrate induced symptoms on onion leaves

We generated a derivative strain of *P*. *stewartii* subsp. *indologenes* PNA14-12 by crossing the plasmid pSC201 into the *halA* gene. This approach aided in bringing the expression of the Halophos cluster under the control of the P*rhaB* promoter. The scheme is depicted in S3 Fig and is similar to the scheme used to express pantaphos used by Polidore et al, 2020 [11]. When cultured with rhamnose supplementation, the filtered culture supernatant of strain 14-12WT pSC201∷*halA* and the strain PNA14-12Δ*pepM* pSC201∷*halA* pBS46∷*pepM* induced necrosis on onion leaves, while the strain PNA14-12Δ*pepM* pSC201∷*halA* did not (Fig 6). When cultured without rhamnose supplementation, the culture supernatant of all three strains did not cause any symptoms (Fig 6).

**Fig 6.**
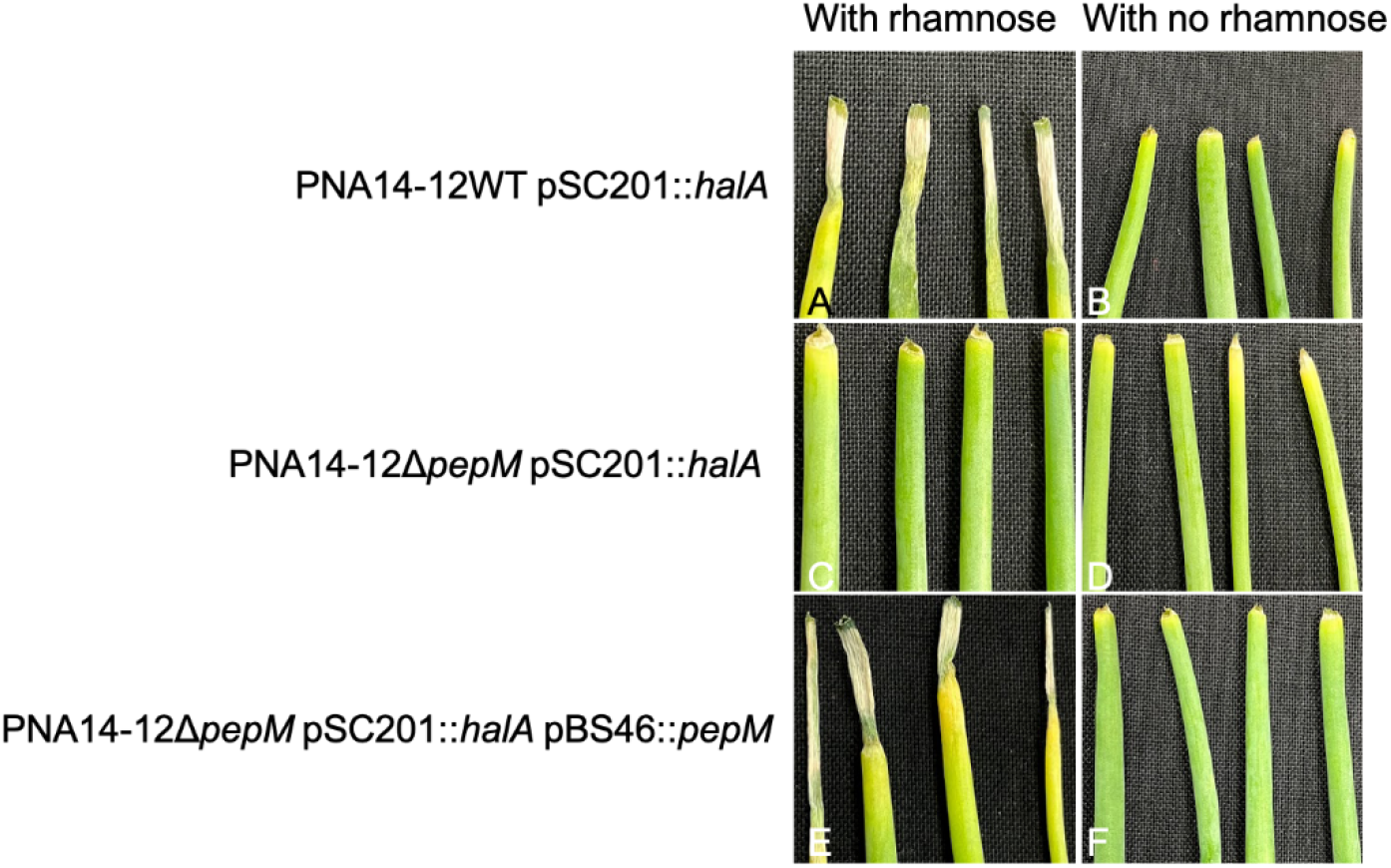
Cell-free culture filtrate induced symptoms on onion leaf. Bacteria were grown in modified Coplin lab medium (mCLM) for 24 h and centrifuged. The supernatant was filtered and 20 μl were inoculated onto cut onion leaf tip. Images were taken 4 days after inoculation. (A) PNA14-12WT pSC201∷*halA* in mCLM + trimethoprim + rhamnose. (B) PNA14-12WT pSC201∷*halA* in mCLM + trimethoprim. (C) PNA14-12Δ*pepM* pSC201∷*halA* in mCLM + trimethoprim + rhamnose. (D) PNA14-12Δ*pepM* pSC201∷*halA* in mCLM + trimethoprim. (E) PNA14-12Δ*pepM* pSC201∷*halA* pBS46∷*pepM* in mCLM + gentamicin + trimethoprim + rhamnose. (F) PNA14-12Δ*pepM* pSC201∷*halA* pBS46∷*pepM* in mCLM + gentamicin + trimethoprim. The experiments were conducted twice with similar results.

### The *hrcC* but not the *pepM* gene is important for the pathogenicity of *P. stewartii* subsp. *indologenes* on millet

On pearl millet leaves, *P. stewartii* subsp. *indologenes* PNA03-3Δ*hrcC* reached significantly lower population levels than PNA03-3 wildtype, 03-3Δ*pepM*, and 03-3Δ*hrcC* pBS46∷*hrcC_03-3_* at 4 dpi (*P* < 0.0001) (Fig 7A). In terms of symptom development, at 4 dpi, 03-3WT, 03-3Δ*pepM*, and 03-3Δ*hrcC* pBS46∷*hrcC_03-3_* developed obvious lesions, while 03-3Δ*hrcC* did not develop any symptoms on inoculated pearl millet leaves (Fig 7B). Similarly, *P. stewartii* subsp. *indologenes* PNA14-12Δ*hrcC* reached significantly lower population levels than PNA14-12 wildtype, 14-12Δ*pepM*, and 14-12Δ*hrcC* pBS46∷*hrcC_03-3_* at 4 dpi (*P* < 0.0001) (Fig 7C). In terms of symptom development, at 4 dpi, 14-12WT, 14-12Δ*pepM*, and 14-12Δ*hrcC* pBS46∷*hrcC_03-3_* developed obvious lesions, while 14-12Δ*hrcC* did not develop any symptoms on inoculated pearl millet leaves (Fig 7D).

**Fig 7.**
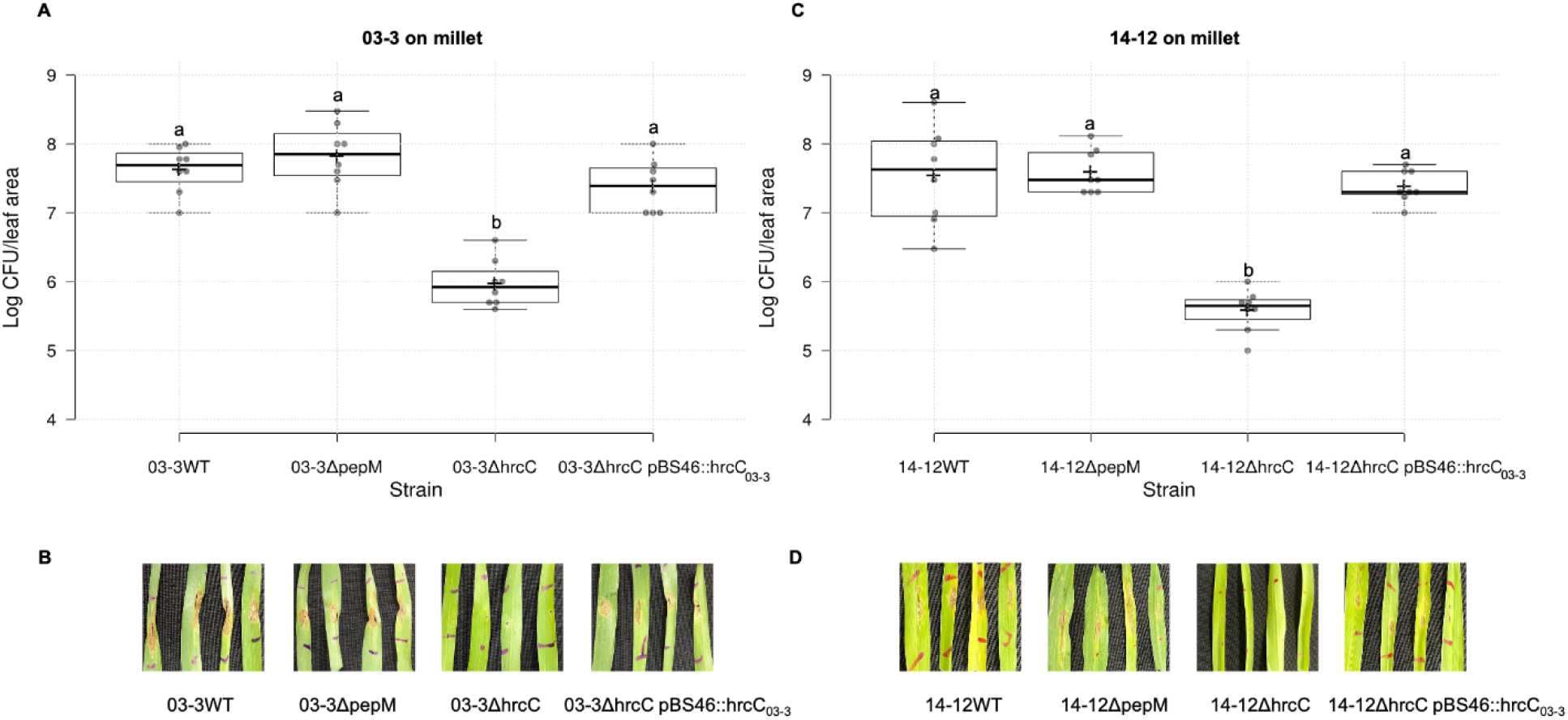
*hrcC* was vital for PNA03-3 and PNA14-12 pathogenicity on pearl millet. Bacterial population levels (A and C) in millet tissues inoculated with *Pantoea stewartii* subsp. *indologenes* PNA03-3 and PNA14-12 wildtype and mutants and representative symptoms produced by 4-week-old millet leaves (B and D). PNA03-3 wildtype (WT), 03-3*ΔpepM* mutant, 03-3*ΔhrcC* mutant, and 03-3Δ*hrcC* pBS46∷*hrcC_03-3_* complement strains, PNA14-12 wildtype (WT), 14-12*ΔpepM* mutant, 14-12*ΔhrcC* mutant, and 14-12Δ*hrcC* pBS46∷*hrcC_03-3_* complement strains were inoculated into millet leaves at 10^6^ CFU/ml. Samples and images were taken at 4 days post inoculation (dpi). Two leaf disks (0.4 cm diameter) were sampled from four leaves, macerated in sterilized H_2_O, and plated on LB agar with rifampicin. The colonies were counted 24 h after incubation and converted to Log_10_ CFU/leaf area (cm^2^). Center lines show the medians; box limits indicate the 25^th^ and 75^th^ percentiles; whiskers extend 1.5 times the interquartile range from the 25^th^ and 75^th^ percentiles; crosses represent sample means; data points are plotted as grey circles as determined by R software. The experiment was conducted twice and all sample points (*n* = 8) were shown. Different letters indicate significant differences (*P* = 0.05) among treatments according to Tukey-Kramer’s honestly significant difference test.

### Importance of *hrcC* and *pepM* genes in tobacco hypersensitive-like cell death response

Hypersensitive response (HR) assays were assessed on tobacco to determine the function of T3SS in *P. stewartii* subsp. *indologenes*. We also evaluated if the products from the HiVir and Halophos clusters in *Pantoea* spp. (*P. stewartii* subsp. *indologenes* and *P. allii*) induced HR-like responses on tobacco. Leaves infiltrated with the suspensions of *P. stewartii* subsp. *indologenes* 03-3WT, 03-3Δ*hrcC*, 03-3Δ*pepM*, 03-3Δ*hrcC* pBS46∷*hrcC*_03-3_, 03-3Δ*pepM*Δ*hrcC* pBS46∷*hrcC*_03-3_, and 03-3Δ*pepM*Δ*hrcC* pBS46∷*pepM*_14-12_ developed an HR-like cell death (CD) response after 48 h (S4A Fig). The double mutant 03-3Δ*pepM*Δ*hrcC* could not induce cell death (S4A Fig). Leaves infiltrated with suspensions of *P. stewartii* subsp. *indologenes* 14-12WT, 14-12Δ*pepM*, 14-12Δ*hrcC* pBS46∷*hrcC*_03-3_, 14-12Δ*pepM*Δ*hrcC* pBS46∷*hrcC*_03-3_ developed an HR-like CD response after 48 h (S4B Fig). Leaves infiltrated with 14-12Δ*hrcC*, 14-12Δ*pepM*Δ*hrcC*, and the control did not induce cell death (S4B Fig). Leaves infiltrated with suspensions of *P. allii* 24248WT, 24248Δ*pepM*, 24248Δ*pepM* pBS46∷*pepM*_24248_, and the control did not induce an HR-like CD response (S4C Fig).

## Discussion

Many plant pathogenic bacteria use secretion systems to deliver proteins to plant cell targets and these secretion systems are often plant pathogenicity determinants [9]. The T3SS was studied extensively in *P. stewartii* subsp. *stewartii* in its interactions with maize [15]. *Pantoea stewartii* subsp. *indologenes* also encodes T3SS, which shows the closest similarity to *P. stewartii* subsp. *stewartii*. However, our study showed that *P. stewartii* subsp. *indologenes* did not require a functional T3SS to infect onion. When *hrcC*, the gene encoding the conserved T3SS outer membrane secretin, was deleted in PNA03-3 and PNA14-12, the *hrcC* mutants still caused onion tissue necrosis (Figs 2 and 3), but lost their pathogenicity on pearl millet and the ability to cause tobacco HR (Figs 7 and S4). This indicated that *P. stewartii* subsp. *indologenes* used alternative virulence strategies to cause disease on onion other than the typical T3SS.

The *pepM* deletion mutants of HiVir (PNA03-3) and Halophos (PNA14-12) clusters did not develop any obvious symptoms on onion leaves and on red onion scales compared to the corresponding wildtype strains (Figs 2 and 3). This indicated that both phosphonate biosynthetic gene clusters are important for the onion pathogenicity for the two *P. stewartii* subsp. *indologenes* strains (PNA14-12 and PNA03-3). The complementation of each gene in the corresponding mutant strains restored the phenotypes. In addition, the *pepM* gene from PNA14-12 could complement 03-3Δ*pepM* as the 03-3Δ*pepM* pBS46∷*pepM_14-12_* restored the onion scale clearing phenotype (Fig 2B). The *in planta* population assays showed that *pepM* mutants grew to significantly lower population levels compared to the wildtype strains. This gene cluster was given the name ‘Halophos’ as the wildtype strain showed a pink halo phenotype when inoculated on red onion scales (Figs 3B, 4B and S3), a phenotype distinctive from the white hard-edged clearing zones observed with *P. ananatis* inoculation on the red onion scale [16].

Two distinct phosphonate clusters (HiVir and Halophos) were responsible for the onion pathogenicity of *P. stewartii* subsp. *indologenes*. Host range expansion of *P. stewartii* subsp. *indologenes* was likely associated with the acquisition of the HiVir or Halophos clusters. The sequence features (GC/Nc/CAI) of the Halophos gene cluster are different from the whole genome features (Table 1), indicating an HGT origin. The presence of the nearby primase gene can also indicate an HGT region [17]. The Nc and CAI values are measures of synonymous codon usage bias. The higher Nc and lower CAI values in the Halophos gene cluster compared to the whole genome of PNA14-12 implies recent acquisition via horizontal gene transfer. In Georgia USA, onion and millet rotation were once recommended and some organic onion growers still follow this rotation. We speculate that *P. stewartii* subsp. *indologenes* might have horizontally acquired virulence factors from environmental or plant-associated microbes associated with millet or other weeds capable of supporting onion-infecting bacteria as well as *P. stewartii* subsp. *indologenes* populations.

Plant pathogens can be classified according to their lifestyles. Biotrophic pathogens require living plant cells to proliferate, whereas necrotrophic pathogens kill plant cells to proliferate. Hemibiotrophic pathogens initially colonize hosts via biotrophic invasion and later switch to necrotrophic growth [18]. Likewise, we speculate that *P. stewartii* subsp. *indologenes* may utilize T3SS, which is an indicator of biotrophic or hemi-biotrophic lifestyle to colonize and infect millets, but on onion it may utilize a different set of pathogenicity factors, HiVir or Halophos, which results in host cell death, an indicative of a necrotrophic lifestyle. These observations potentially suggest that onion-pathogenic *P. stewartii* subsp. *indologenes* may utilize hemi-biotrophic or necrotrophic lifestyles preferentially when they are present on millet vs. on onion. Future detailed studies may shed some light on this dual lifestyle and its importance in host-range expansion.

Similar to PNA14-12, *P. allii* LMG24248 also showed a pink halo phenotype on the red onion scale, while its *pepM* mutant did not develop any symptoms (Fig 4). This indicated that *pepM* of the Halophos gene cluster was also important for onion pathogenicity in *P. allii*, which has never been demonstrated experimentally earlier. We also note that *P. allii* strains do not possess T3SS; however, it was present in *P. stewartii* subsp. *indologenes*.

In this study, we showed that *pepM* was essential for causing necrosis on both onion leaves and scales in two *Pantoea* spp*., P. stewartii* subsp. *indologenes* and *P. allii*. Furthermore, Halophos-like gene clusters were also present in strains belonging to the genera *Erwinia, Pseudomonas, Xenorhabdus*, and *Photorhabdus. Xenorhabdus* and *Photorhabdus* are nematode and arthropods symbionts. Interestingly, a HiVir-like cluster was also found in *Photorhabdus* [11]. Many *Erwinia* and *Pseudomonas* species are common plant pathogens on various crops [1, 19, 20]. Despite their presence in various bacterial genera, it would be interesting to test if Halophos toxins are host-specific, and their interactions with insects or nematodes. It is interesting to note that PNA03-3Δ*hrcC* caused an HR-like CD response but PNA03-3Δ*pepM*Δ*hrcC*, PNA14-12Δ*hrcC*, and *P. allii* LMG24248WT did not, suggesting the HiVir toxin from PNA03-3 but not the Halophos toxins from PNA14-12 and *P. allii* LMG24248 is toxic to tobacco leaf (S4 Fig). This also indicates the modes of action between HiVir and Halophos products are different.

Halophos was predicted *in silico* to encode a putative phosphonate biosynthetic gene cluster. In this study, we characterized individual genes in the Halophos gene cluster, and found that *halB-H* but not *halA, halI, halJ, halK* genes contribute to onion pathogenicity (Fig 5). We showed that by transferring a minimal cluster of *halB to halH* to an onion-non-pathogenic strain PNA15-2, the strain was able to cause foliar necrosis on onion (Figs 5D and 5E). However, the mean lesion length of PNA15-2 pBS46∷*halB-H* was relatively small. This may be due to the low expression level of the plasmid carrying a large fragment of *halB to halH* (8,614 bp) or other components present in PNA14-12 but absent in PNA15-2 that might be needed to facilitate the expression of *halB to halH*. In addition, to our surprise, *halI*, the MFS transporter gene from the Halophos cluster did not contribute to onion pathogenicity (Fig 5), while *hvrI*, the MFS transporter gene from the HiVir cluster in *P. ananatis* was shown to be important for onion leaf virulence [8]. We speculate that PNA14-12 does not rely on *halI* but other unknown means to transport its product of Halophos out of the cell.

On the other hand, only *pepM* and the MFS transporter gene from the HiVir cluster were previously characterized and shown to be important for onion pathogenicity [7, 8, 11]. The individual contributions of other genes in the HiVir cluster remain to be determined. Identifying the minimal number of genes required for phosphonate biosynthesis may help in the identification of key players involved in the biosynthetic pathway and may aid in understanding how phosphonates are synthesized. The phosphonate structure of the HiVir gene cluster was recently determined as 2-(hydroxy[phosphono]methyl)maleate [11]; however, the products of the predicted phosphonate biosynthetic cluster ‘Halophos’ and their biological functions are currently unknown. An unknown thiotemplated cluster type (S5 Fig) was predicted by the PRISM program (version 4.4.5) [21] based on *halE/pepM gene* and *halJ* gene from the PNA14-12 Halophos gene cluster. However, the predicted structure did not incorporate the function of the other genes in the Halophos cluster. Understanding the products, the regulation of phosphonate biosynthesis, and the synthetic stages may help provide targets for developing specific enzyme inhibitors against plant-pathogenic *Pantoea* carrying Halophos.

In conclusion, a unique gene cluster Halophos, responsible for onion tissue necrosis was found in *P. allii* and *P. stewartii* subsp. *indologenes*. Future research on Halophos structure and its target are needed for developing strategies to manage *Pantoea* spp. in onion.

## Materials and Methods

### Bacterial strains and inoculum preparation

Bacterial strains and plasmids used in the study are listed in S2 Table. Naturally occurring, rifampicin-resistant strains of representative *Pantoea* strains were selected. *Pantoea* strains were routinely cultured on nutrient agar at 28°C for 1 day and *Escherichia coli* strains on Luria-Bertani broth or agar at 37°C for 1 day. When required, media were supplemented with kanamycin at 50 μg/ml, trimethoprim at 100 μg/ml, gentamycin at 20 μg/ml, X-gluc at 60 μg/ml, rifampicin at 50 μg/ml, and diaminopimelic acid (DAP) for *E. coli* RHO5 at 300 μg/ml. To prepare *Pantoea* inocula, strains were cultured in LB at 28°C in a rotary shaker at 200 rpm for approximately 16 h. Subsequently, the cultures were centrifuged at 16,100 x g for 1 min and the supernatants were decanted. The resulting pellets were resuspended in sterilized distilled water (sdH_2_O). The bacterial concentrations were then adjusted to an optical density of 0.3 at 600 nm [~ 10^8^ colony forming units (CFU/ml)] using a spectrophotometer, and adjusted to the desired concentration by 10-fold serial dilutions in sdH_2_O.

### Mutant constructions

To determine the role of *pepM* in *Pantoea* pathogenicity, unmarked deletions of *pepM* gene in *Pantoea allii* LMG 24248, and *P. stewartii* subsp. *indologenes* PNA03-3 and PNA14-12 were made using the pR6KT2GW allelic exchange vector following the method described in [7], and named 24248Δ*pepM*, PNA03-3Δ*pepM*, and PNA14-12Δ*pepM*, respectively. The primers used are listed in S3 Table. dsDNA containing the *pepM* flanking sequences were synthesized (Twist Biosciences, San Francisco, CA) and listed in S4 Table. Mutants were confirmed by PCR and Sanger sequencing. To determine the role of *hrcC* in *P. stewartii* subsp. *indologenes* virulence, PNA03-3Δ*hrcC* and PNA14-12Δ*hrcC* were made. Double mutants of *pepM* and *hrcC* were made using the Δ*pepM* mutant for conjugation following the deletion procedure as described above.

For the complementation, the *hrcC* gene sequence from PNA03-3 and *pepM* gene sequences from both PNA14-12 and LMG24248 including their native ribosomal binding sites were PCR amplified and used to create pBS46∷*hrcC_03-3_/pepM_14-12_/pepM_24248_* constructs for conjugation with corresponding deletion mutants following the method described in [7].

To determine the role of *hal* genes from the Halophos cluster, unmarked deletions of *halA, halB, halC, halD, halF, halG, halH, halI, halJ*, and *halK* genes in PNA14-12 were made using the pR6KT2GW allelic exchange vector following the deletion procedure as described above. The complementations were made as described above for *halB, halC, halD, halF, halG*, and *halH* single mutants. In addition, we constructed a plasmid containing sequence from *halB to halH* and transformed this plasmid pBS46∷*halB-H* into an onion-non-pathogenic *P. stewartii* subsp. *indologenes* strain PNA15-2.

### *In planta* population and symptomology on onion caused by select *Pantoea* strains

*Pantoea* strains (wildtypes, mutants and their corresponding complemented strains) were inoculated on leaves and red onion scale as previously described [16]. Briefly, red onion bulbs (cv. Red Barret) were sliced into small square slices (approximately 2 cm x 2 cm), sterilized in 0.5% sodium hypochlorite, and washed with tap water. A 10 μl pipette tip was used to penetrate the onion scale at the center with finger pressure. Ten microliters of a 1 x 10^8^ CFU/ml bacterial suspension (1×10^6^ CFU) were deposited at the wounded area. The sdH_2_O was used as the negative control. Samples and images were taken at 4 days post inoculation (dpi). Scale tissue samples (approximately 0.2 cm x 0.2 cm) were cut using a sterile blade 0.5 cm away from the inoculation point, weighed, and placed in 2 ml tubes containing three sterile glass beads (3 mm) and 500 μl sdH_2_O. Tissue samples were macerated in a bead mill homogenizer (Omni International Inc, Kennesaw, GA, USA) four times for 30 seconds each at 4 m/s speed. A ten-fold dilution series of the macerates was made in sterile sdH_2_O to 10^−6^ and each dilution was plated as 10 μl droplets on LB agar with rifampicin. Colonies were counted 24 h after incubation and converted to Log_10_ CFU/g. Four replicates per strain were used for one experiment and the experiment was conducted at least twice.

Foliage inoculation assays were conducted as described by Koirala *et al*. (2021) [13]. Briefly, onion (cv. Century) seedlings were established in plastic pots and maintained in a greenhouse at 25-30°C. Onion plants about six- to eight-week-old were inoculated after cutting the leaf 1 cm from the apex with a pair of scissors sterilized with 70% ethanol. Using a micropipette, 10 μl of 1 x 10^6^ CFU/ml bacterial suspension (~ 1 x10^4^ CFU/leaf) was deposited diagonally opposite to each other at the cut end of the leaf twice. Seedlings inoculated with sdH_2_O as described above were used as negative controls. The strains used in the inoculation included PNA14-12 wildtype, PNA03-3 wildtype, LMG24248, and their corresponding mutants and complements are listed in S2 Table. At 4 dpi, lesion lengths of the inoculated leaf areas were measured. Leaf tissues (0.5 cm long) were taken 0.3 cm away from the inoculation point, processed similarly to the scale samples, and the enumerated colonies were converted to Log_10_ CFU/leaf area (cm^2^).

### Symptomology on onion caused by select *Pantoea* strains

The following strains, South Africa *P. allii* strains, single mutants of *hal* genes and *hal* complement strains, PNA14-12, PNA15-2, PNA15-2 pBS46∷*halB-H*, were inoculated as described above to compare their symptoms on onion. For both red onion scale assays and foliage inoculation assays, 10 μl of 1 x 10^6^ CFU/ml bacterial suspensions were used. The pictures and lesion lengths were recorded at 5 dpi.

### Symptomology on onion leaf caused by culture filtrates of *Pantoea* strains

We hypothesized that the Halophos gene cluster in *P. stewartii* subsp. *indologenes* PNA14-12 produced a secondary metabolite product, and the product alone could cause symptoms on onion leaves. A single crossover approach similar to that used by Polidore *et al*. 2021 [11] for *in vitro* induction of HiVir was used to drive the expression of the Halophos cluster from the rhamnose inducible P*rhaB* promoter by recombining plasmid pSC201∷*halA* before the *halA* gene of the Halophos. The scheme was depicted in S3 Fig. The first gene *halA* in the Halophos gene cluster from PNA14-12 including 39 bp before the start codon was PCR amplified, purified, and mixed with the plasmid pSC201 (cut with *XbalI* and *SphI*) in a Gibson-assembly mix (NEB). The reaction was transformed into *E. coli* DH5α, and plated on LB agar with trimethoprim. The resulting plasmid was confirmed by sequencing, and transformed into *E. coli* RHO5. Bi-parental conjugation was performed by plating a mixture of *E. coli* strain RHO5 pSC201∷*halA* with the PNA14-12 wildtype, PNA14-12Δ*pepM*, and PNA14-12Δ*pepM* pBS46∷*pepM* on LB agar with DAP. The mixed cultures were streaked onto LB agar with trimethoprim. The single colonies were streaked on LB agar with trimethoprim and stored. The chromosomal insertion of the plasmid was confirmed by PCR (S3 Table).

Bacteria were grown in the modified Coplin lab medium (mCLM) [8] at 28°C for 24 h with shaking at 200 rpm. In order to maintain plasmid insertion, the medium was amended with trimethoprim. For the complemented strain, the medium was also amended with gentamicin. All strains were grown in 5 ml of this broth with and without 0.5% rhamnose. The cultures were centrifuged and the supernatants were filter-sterilized through a 0.2-μm filter to obtain a crude culture filtrate. Before use, 100 μl of this crude culture filtrate was plated on the nutrient agar medium in two replicates to confirm if it was devoid of any bacterial contamination. No colonies were grown after two days of incubation at 28°C. Twenty microliters of the crude culture filtrates were inoculated onto the cut end of onion leaf tips as described previously. Inoculated seedlings were assessed for foliar necrosis at 4 dpi and subsequently images were taken. The experiments were conducted twice.

### *In planta* population and symptomology on pearl millet caused by *Pantoea* strains

We hypothesized that *hrcC* but not *pepM* was important for pearl millet pathogenicity. To determine the role of *pepM* and *hrcC* in the pathogenicity of *P. stewartii* subsp. *indologenes* on pearl millet, *Pantoea* strains were inoculated on 4-week-old pearl millet seedlings (*Pennisetum glaucum* cv. TifGrain 102) at 1 x 10^6^ CFU/ml by syringe infiltration. Pearl millet seedlings were established under the same growth condition as described above for onion seedlings. Symptoms were documented on the inoculated leaves, by taking photographs at 4 dpi. *In planta* populations of inoculated bacterial strains were assessed at 4 dpi by excising two leaf discs per replicate using a 0.4 cm diameter cork-borer. Four replicates were sampled per strain and the experiments were conducted twice. Samples were macerated in sdH_2_O and plated on LB agar with rifampicin. Colonies were counted 24 h after incubation and converted to Log_10_ CFU/leaf area (cm^2^).

Analysis of variance (ANOVA) was conducted on Log_10_ CFU/leaf area (cm^2^), Log_10_ CFU/g, and lesion length data using JMP statistical analysis software (version Pro 16; SAS Institute Inc., Cary, NC). The effect of strain on the data collected was compared using the Tukey-Kramer’s honestly significant difference (HSD) test. The boxplot was generated using the BoxPlotR web tool [22].

### Tobacco infiltration assay

*Pantoea* spp. strains were grown overnight in mCLM [8] at 28°C with shaking at 200 rpm. The following day, overnight cultures were infiltrated into tobacco leaf panels using a plastic needleless syringe. The infiltrated areas were outlined with a black permanent marker. Sterile mCLM was used as a control. Leaf panels were observed after 48 h for hypersensitive-like cell death responses. The assay was performed on two or three leaves from different plants and was conducted three times.

### Identification and analysis of homologous Halophos biosynthetic gene clusters in bacteria

To assess whether the Halophos gene cluster was introduced by horizontal gene transfer, genomic islands in PNA14-12 were predicted using IslandViewer 4 [23]. We also computed the guanine-cytosine (GC) content, the effective number of codons (Nc), and the codon adaptation index (CAI) using CodonW Version 1.4.4 [24]. In addition, NCBI blastn and tblastn were used to identify bacterial genomes that contain Halophos-like gene clusters. First, the nucleotide sequence of the Halophos gene cluster from PNA14-12 was used to search the NCBI Nucleotide collection database. After identifying Halophos-like gene clusters in several genera, the nucleotide sequence was searched against the NCBI Whole-genome shotgun contigs database, by specifying genera *Pantoea, Pseudomonas, Erwinia, Xenorhabdus*, and *Photorhabdus* in the search options. The query coverage and percent identity of the blastn searches were recorded.

In order to show the gene synteny of Halophos-like gene clusters, the PepM protein sequence from PNA14-12 was used as the query in the NCBI blastp search. The NCBI protein accessions of the PepM protein sequences that were in the Halophos-like gene clusters were downloaded, and used as the query in the WebFlaGs web tool [25]. Results from representative Halophos types of different species were illustrated in PowerPoint along with a phylogenetic tree based on the corresponding selected *PepM* protein sequences. Ten PepM protein sequences and *Escherichia coli* 2-methylisocitrate lyase sequence (as an outgroup) were used for multiple sequence alignment using MAFFT v7.450 [26] in Geneious Prime^®^ 2021.1.1. The alignment was trimmed to the same length and used for constructing a neighbor-joining tree using the Jukes-Cantor model [27]. The bootstrap support values were calculated using 1,000 replicates.

## Data availability statement

Confirm whether all data reported in the manuscript are publicly available. PLOS requires that authors deposit all reported data and related metadata underlying the study findings in an appropriate public repository unless already provided in the submission.

## Supporting information captions

**S1 Fig. Lesion length of onion leaf**. Onion leaves were inoculated with wildtype and mutants of *Pantoea stewartii* subsp. *indologenes* PNA03-3 (A) and PNA14-12 (B), and *P. allii* LMG24248 (C). Strains were inoculated on the cut end of the onion leaf tip at 10^4^ CFU/leaf. Four replicates per strain were used for one experiment. Lesion lengths were measured at 4 days post inoculation. Center lines show the medians; box limits indicate the 25^th^ and 75^th^ percentiles; whiskers extend 1.5 times the interquartile range from the 25^th^ and 75^th^ percentiles; crosses represent sample means; data points are plotted as grey circles as determined by R software. The experiment was conducted at least three times and all sample points were shown (*n* = 12, 40, and 16 for graph A, B, C, respectively). Different letters indicate significant differences (*P* = 0.05) among treatments according to Tukey-Kramer’s honestly significant difference test.

**S2 Fig. Representative symptoms produced on onion scales and leaves inoculated with *Pantoea allii*.** *Pantoea allii* strains were inoculated onto the red onion scales and leaves at 10^6^ CFU. The experiments were conducted twice with similar results.

**S3 Fig. Steps involved in Halophos expression induction with rhamnose**. (A) Plasmid pSC201∷*halA*, which carries the *halA* gene under the control of the rhamnose-inducible P*rhaB* promoter, was introduced to PNA14-12 wildtype, PNA14-12Δ*pepM*, and PNA14-12Δ*pepM* pBS46∷*pepM* via biparental conjugation. (B) The graph (not to scale) showed the chromosomal insertion of the plasmid pSC201∷*halA* after homologous recombination. The resulting strains, maintained by growing in trimethoprim, expressed the entire Halophos operon from the P*rhaB* promoter.

**S4 Fig. Tobacco Hypersensitive Response (HR) like Cell Death (CD) response to *Pantoea* strains**. (A) *Pantoea stewartii* subsp. i*ndologenes* 03-3WT, mutants, and selected complement strains. (B) *P. stewartii* subsp. *indologenes* 14-12WT, mutants, and selected complement strains (C) *P. allii* 24248WT, mutant, and selected complement strains. All strains were grown overnight in mCLM at 28°C with shaking. The overnight culture was used to infiltrate panels of tobacco leaves with a blunt end syringe. Sterile mCLM was used as a control. Tobacco panels were evaluated 48 h post inoculation. Representative images of the HR-like CD response are presented next to our interpretation of the result. The experiment was conducted three times.

**S5 Fig. The predicted structure of an unknown thiotemplated cluster type associated with Halophos**. The structure was predicted by the PRISM program based on *halE/pepM gene* and *halJ* gene from the PNA14-12 Halophos gene cluster. The predicted structure did not incorporate the function of the other genes in the Halophos cluster.

**S1 Table. Strains carrying Halophos-like gene clusters identified in the GenBank databases.**

**S2 Table. Strains and plasmids used in this study. S3 Table. Oligonucleotide primers used in this study.**

**S4 Table. Synthesized dsDNA synthesized by Twist Biosciences for making deletion constructs.**

